# Why evolve reliance on the microbiome for timing of ontogeny?

**DOI:** 10.1101/665182

**Authors:** C. Jessica E. Metcalf, Lucas P. Henry, María Rebolleda-Gómez, Britt Koskella

**Author notes:** Data accessibility*: this paper includes no new data.

## Abstract

The timing of life history events has important fitness consequences. Since the 1950s, researchers have combined first principles and data to predict the optimal timing of life history transitions. Recently, a striking mystery has emerged. Such transitions can be shaped by a completely different branch of the tree of life: bacterial species in the microbiome. Probing these interactions using testable predictions from evolutionary theory could illuminate whether and how host-microbiome integrated life histories can evolve and be maintained. Beyond advancing fundamental science, this research program could yield important applications. In an age of microbiome engineering, understanding the contexts that lead to microbiota signaling shaping ontogeny could offer novel mechanisms for manipulations to increase yield in agriculture, or reduce pathogen transmission by affecting vector efficiency. We combine theory and evidence to illuminate the essential questions underlying the existence of <u>M</u>icrobiome <u>D</u>ependent <u>O</u>ntogenetic <u>T</u>iming (MiDOT) to fuel research on this emerging topic.

## Introduction

Many surprises have emerged as a result of our expanding knowledge of the microbiome, the community of microbial organisms that live in and on eukaryotes. Among the oddest phenomena observed is a defining role of the microbiome in many host species’ ontogeny. Key life-cycle transitions including metamorphosis in mosquitoes [1], maturation in flies [2], and flowering in plants [3] have been tied to the presence or activity of particular microbiota. This observation has been made across a wide array of ontogenetic transitions and taxa (Table 1). Manifestations range from absolute effects (life history transitions that fail to occur in the absence of microbe species), through to continuous modulations (developmental accelerations or decelerations). For example, mosquito larvae will not pupate in the absence of bacteria [1,4], and adding bacteria or yeast rescues development [5]. As an example of modulation, *Drosophila* colonized with *Acetobacter* develop more rapidly than when colonized with *Lactobacillus* [6,7]; and in Brassica, soil microbes associated with drought led to accelerated flowering time compared to microbes associated with wet soils [8]. The question then emerges: why would such an important aspect of the fitness of eukaryotic hosts be driven by the presence or absence of a completely different branch of the tree of life?

**Table 1:**
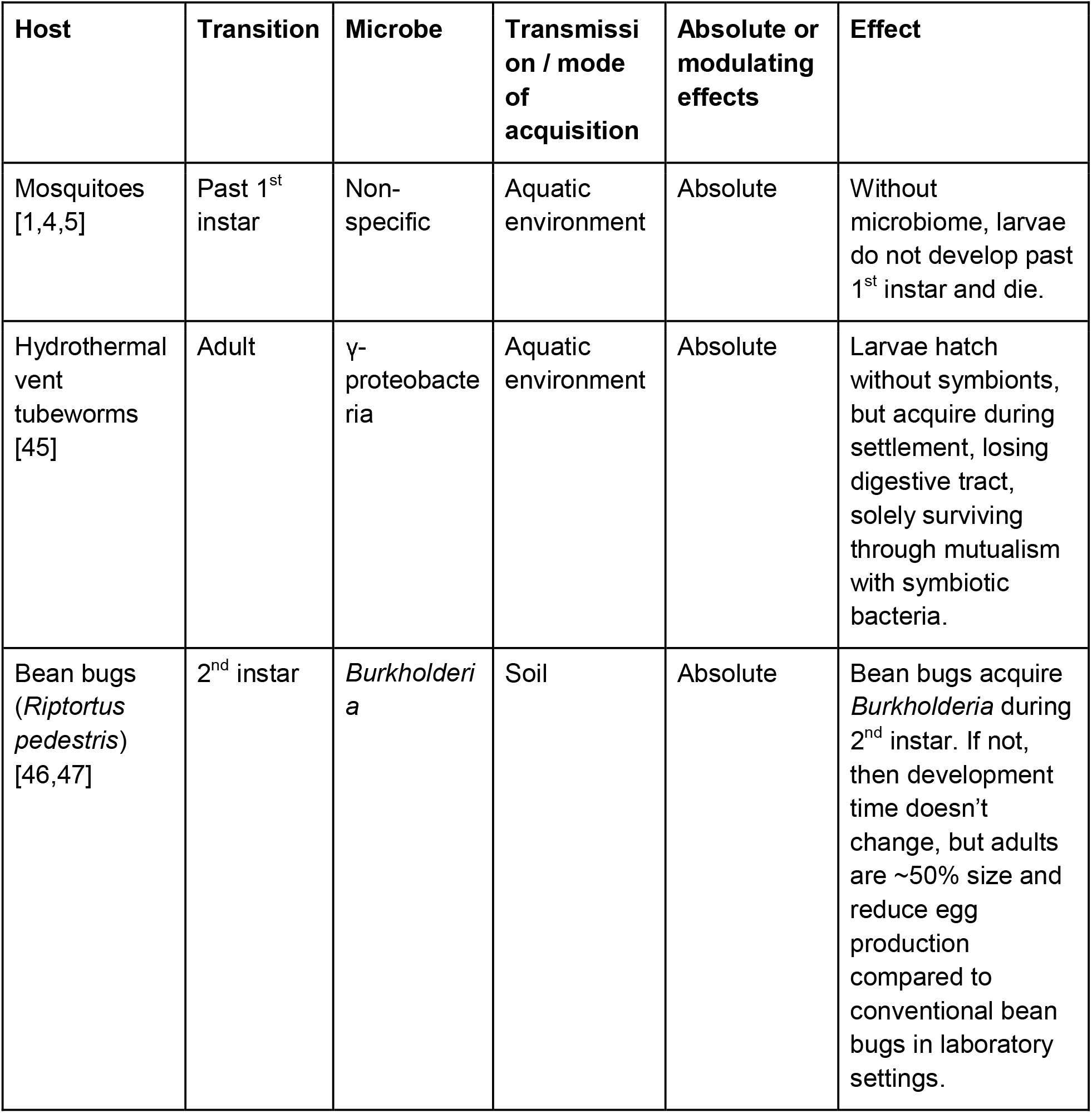
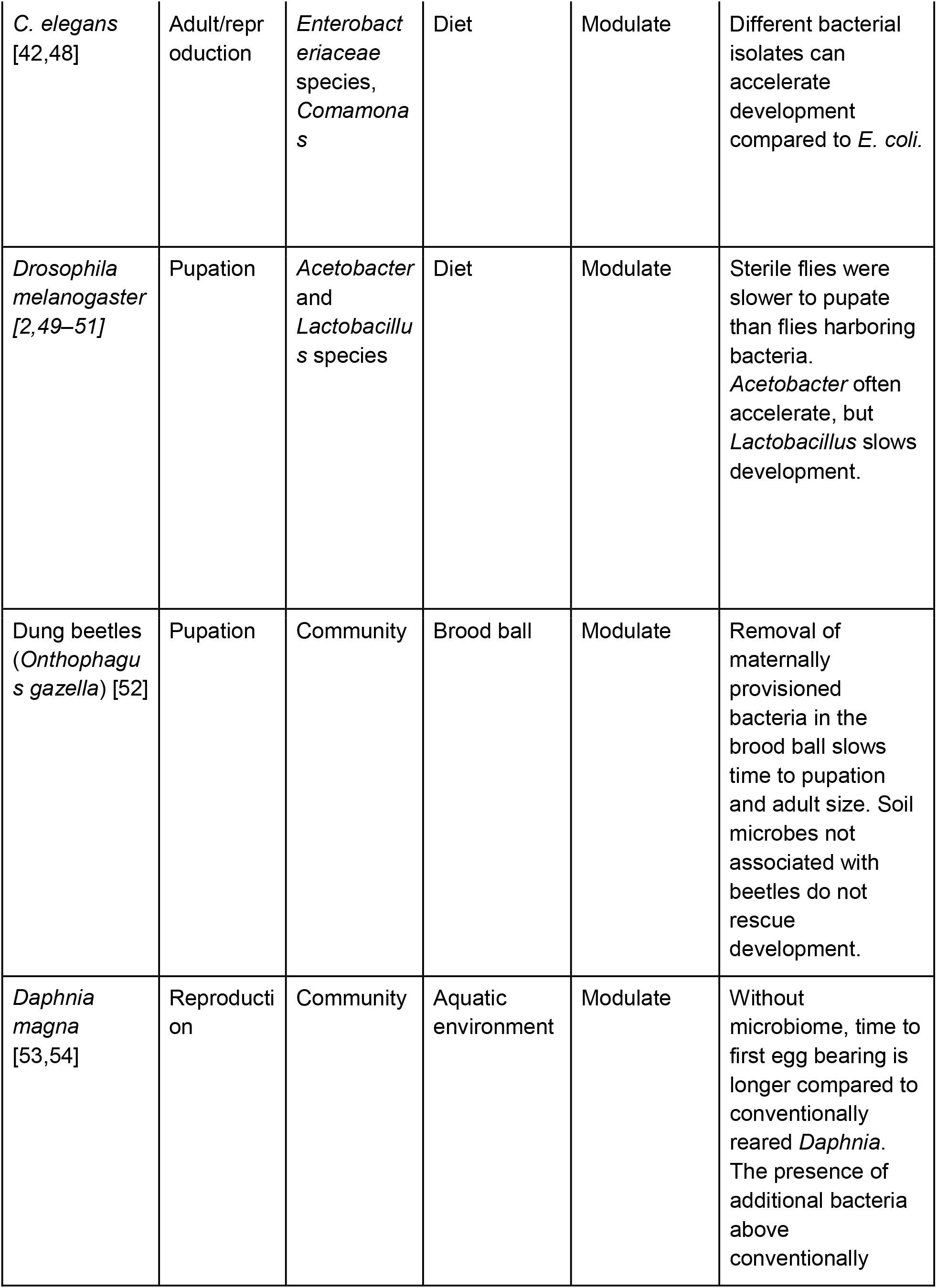
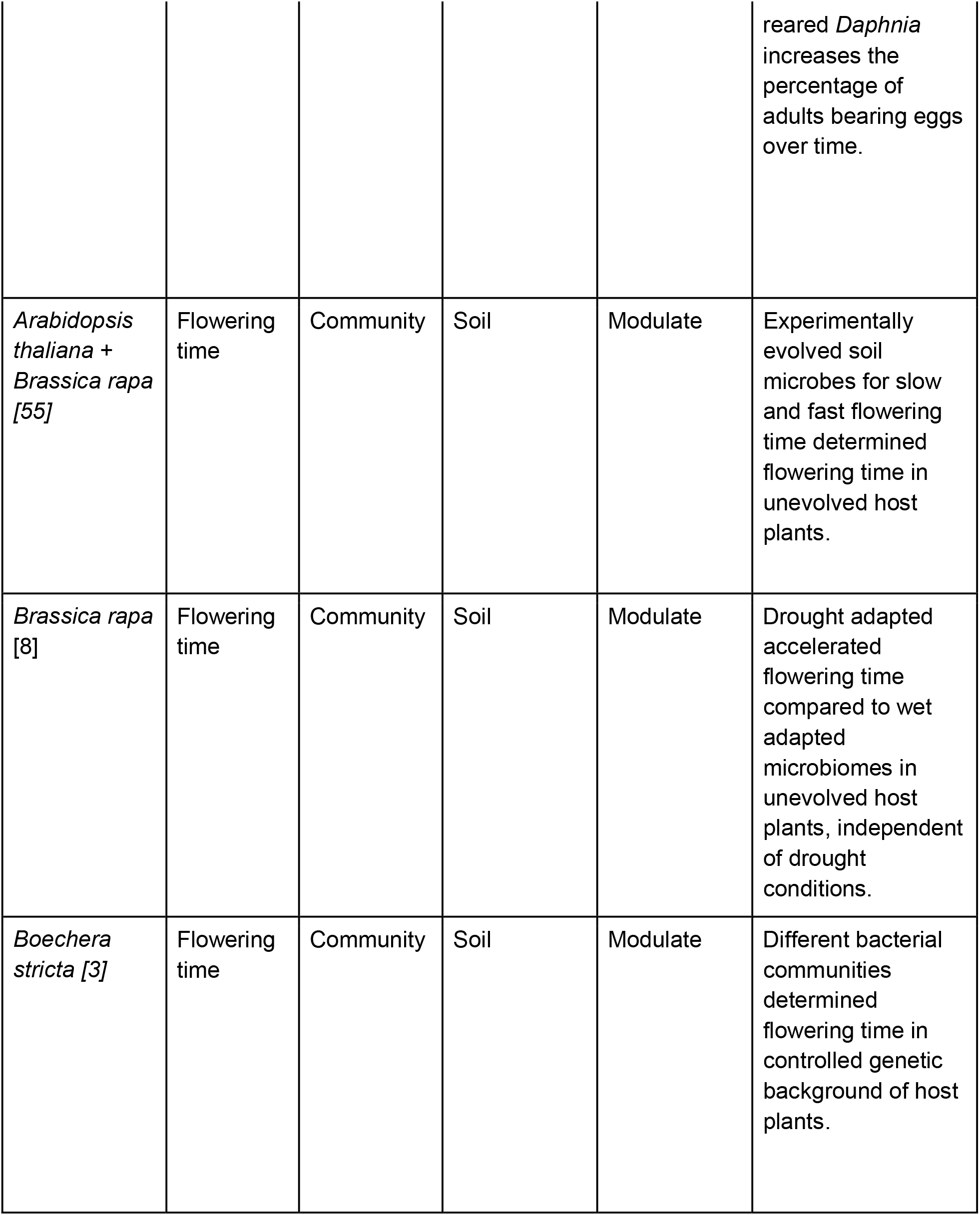
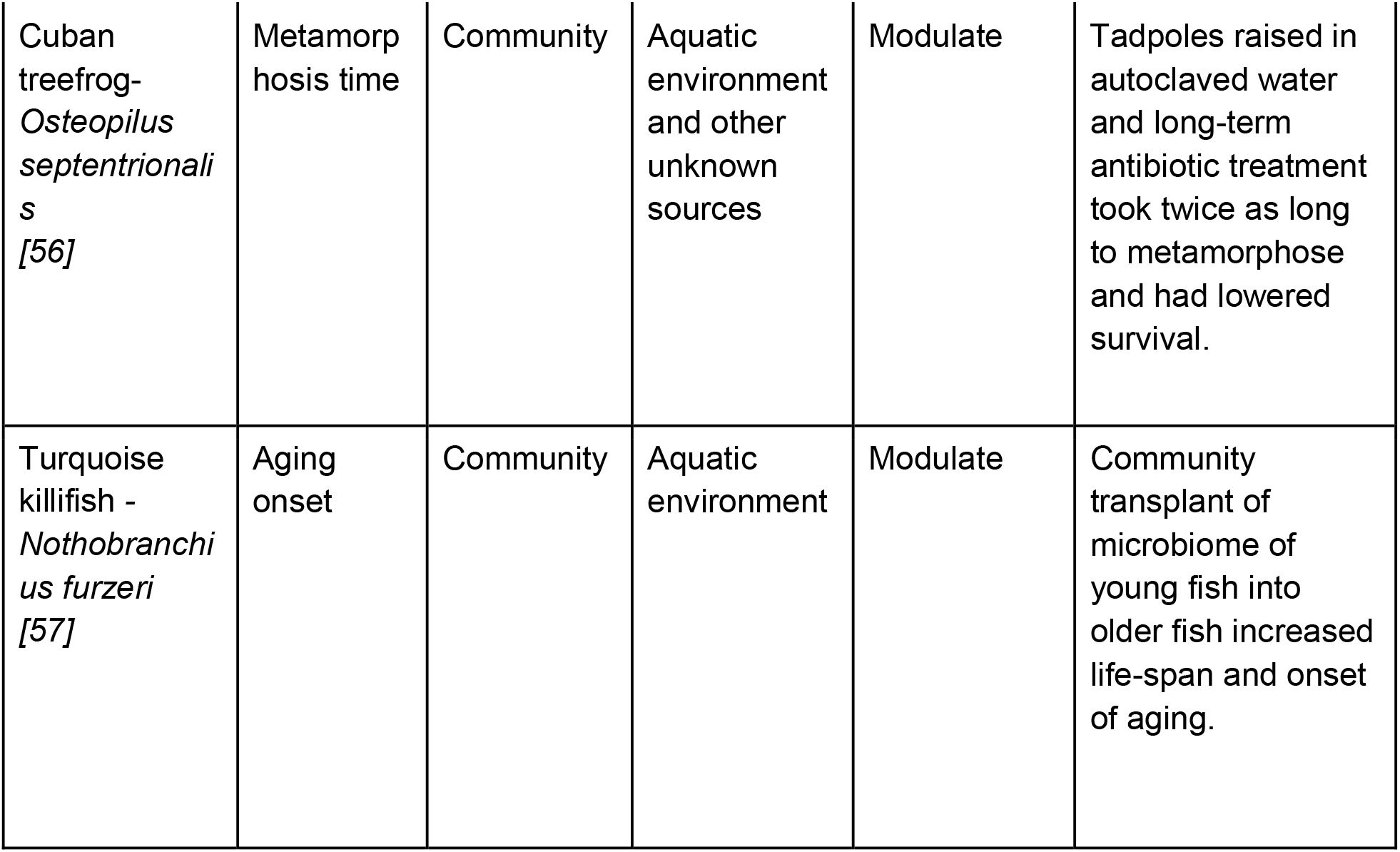
**<u>M</u>icrobiome <u>D</u>ependent <u>O</u>ntogenetic <u>T</u>iming (MiDOT) examples** from host-microbe associations across terrestrial and aquatic organisms. In these examples, experiments controlling host and microbiome variation indicate that the microbiome is a key driver in ontogenetic timing for these environmentally acquired associations between host and microbe. Effects can be absolute (where transition fails to occur in the absence) or modulating (where microbes speed or slow transition).

| Host | Transition | Microbe | Transmission / mode of acquisition | Absolute or modulating effects | Effect |
| --- | --- | --- | --- | --- | --- |
| Mosquitoes [1,4,5] | Past 1<sup>st</sup> instar | Non-specific | Aquatic environment | Absolute | Without microbiome, larvae do not develop past 1<sup>st</sup> instar and die. |
| Hydrothermal vent tubeworms [45] | Adult | γ-proteobacteria | Aquatic environment | Absolute | Larvae hatch without symbionts, but acquire during settlement, losing digestive tract, solely surviving through mutualism with symbiotic bacteria. |
| Bean bugs (*Riptortus pedestris*) [46,47] | 2<sup>nd</sup> instar | *Burkholderia* | Soil | Absolute | Bean bugs acquire *Burkholderia* during 2<sup>nd</sup> instar. If not, then development time doesn't change, but adults are ~50% size and reduce egg production compared to conventional bean bugs in laboratory settings. |
| *C. elegans*<br>[42,48] | Adult/reproduction | *Enterobacteriaceae* species,<br>*Comamonas* | Diet | Modulate | Different bacterial isolates can accelerate development compared to *E. coli*. |
| *Drosophila melanogaster*<br>[2,49–51] | Pupation | *Acetobacter* and<br>*Lactobacillus* species | Diet | Modulate | Sterile flies were slower to pupate than flies harboring bacteria.<br>*Acetobacter* often accelerate, but *Lactobacillus* slows development. |
| Dung beetles (*Onthophagus gazella*) [52] | Pupation | Community | Brood ball | Modulate | Removal of maternally provisioned bacteria in the brood ball slows time to pupation and adult size. Soil microbes not associated with beetles do not rescue development. |
| *Daphnia magna*<br>[53,54] | Reproduction | Community | Aquatic environment | Modulate | Without microbiome, time to first egg bearing is longer compared to conventionally reared *Daphnia*. The presence of additional bacteria above conventionally reared *Daphnia* increases the percentage of adults bearing eggs over time. |
| *Arabidopsis thaliana* + *Brassica rapa* [55] | Flowering time | Community | Soil | Modulate | Experimentally evolved soil microbes for slow and fast flowering time determined flowering time in unevolved host plants. |
| *Brassica rapa* [8] | Flowering time | Community | Soil | Modulate | Drought adapted accelerated flowering time compared to wet adapted microbiomes in unevolved host plants, independent of drought conditions. |
| *Boechera stricta* [3] | Flowering time | Community | Soil | Modulate | Different bacterial communities determined flowering time in controlled genetic background of host plants. |
| Cuban treefrog-<br>*Osteopilus septentrionalis*<br>[56] | Metamorphosis time | Community | Aquatic environment and other unknown sources | Modulate | Tadpoles raised in autoclaved water and long-term antibiotic treatment took twice as long to metamorphose and had lowered survival. |
| Turquoise killifish -<br>*Nothobranchius furzeri*<br>[57] | Aging onset | Community | Aquatic environment | Modulate | Community transplant of microbiome of young fish into older fish increased life-span and onset of aging. |

For such an association to evolve, selection might have acted on the host, on the microbes, or on both. Microbes have clear incentives to manipulate host ontogeny, for example delaying transitions that result in a life stage that curtails their persistence or transmission. Host castration by parasites provides a classic and extreme example: host resources are then diverted away from host reproduction and towards parasite growth [9]. In the other direction, induction of earlier flowering of *Silene viscera* by the anther smut pathogen (*Microbotryum violaceum*) increases the probability of transmission to a new host [10].

A rich array of microbes are known to have adapted to manipulate their host’s ontogeny in these ways. Yet, the other possibility, host adaptation to cue into microbial signaling to determine timing of key transitions, is rarely considered, presumably in part because microbial generation times are so much faster than their hosts. Indeed, it has been suggested that host responses to microbiome species generally reflect the outcome of chance alone [11]. Yet, there is clear theoretical scope for selection on hosts. Empirical support is also growing. For example, shifts in timing of flowering in Brassica associated with microbe presence were found to be associated with host fitness benefits [8]. We refer to this phenomenon as <u>M</u>icrobiome <u>D</u>ependent <u>O</u>ntogenetic <u>T</u>iming (MiDOT).

What circumstances might lead host species to rely on cues from their microbiome to trigger important life history transitions (MiDOT), rather than cues directly reflecting their own internal state or cues gleaned from the abiotic environment? Can we leverage existing tools to go beyond the correlative (host ontogeny in/decreases in the presence of particular microbes) to the predictive (hosts should respond by in/decreasing ontogeny in the presence of particular microbes to maximize fitness) in evaluating the role of MiDOT? Here, we describe relevant results from life history theory; outline how infectious disease ecology informs expectations for patterns of microbiome acquisition; and synthesize these threads to discuss what contexts might allow MiDOT.

### Predictions from life history theory

Life history theory defines how selection shapes the timing of developmental transitions (maturation, eclosion, flowering…) to maximize fitness [12]. A first important result is that delaying life-history transitions is never optimal unless some benefit accrues during the delay, both because of the risk of death over the course of the delay, but also because any delay slows population growth. Beyond this, theory indicates that the optimal delay until a life history transition occurs may hinge either on endogenous features of the host (e.g., growth rate, where reproductive output is size dependent [13]), or exogenous features (e.g., temperature, time of year, where these are associated with better survival of the next life stage), or both.

The timing of reproduction of monocarpic species (where reproduction is fatal) provides a useful template for considering selection pressures on ontogenetic transitions: intersection between theory and empiricism has uncovered important ways in which the heterogeneity of realistic systems modulates theoretical predictions [14,15]. Broadly, theory indicates that monocarpic species should reproduce when the risk of mortality over the next year outweighs the benefits accrued by delaying and growing (since larger size is associated with greater seed production [13]; Figure 1). Such dependence on growth rate highlights that it is important to distinguish between scenarios where microbes are only associated with ontogenetic timing, vs. scenarios where microbes are associated with timing, but also other fitness features (e.g., rate of growth [2,16], survival). Such effects have important implications for what the presence or absence of particular microbial species indicates relative to the optimal timing of flowering (see below, Table 2).

**Figure 1:**
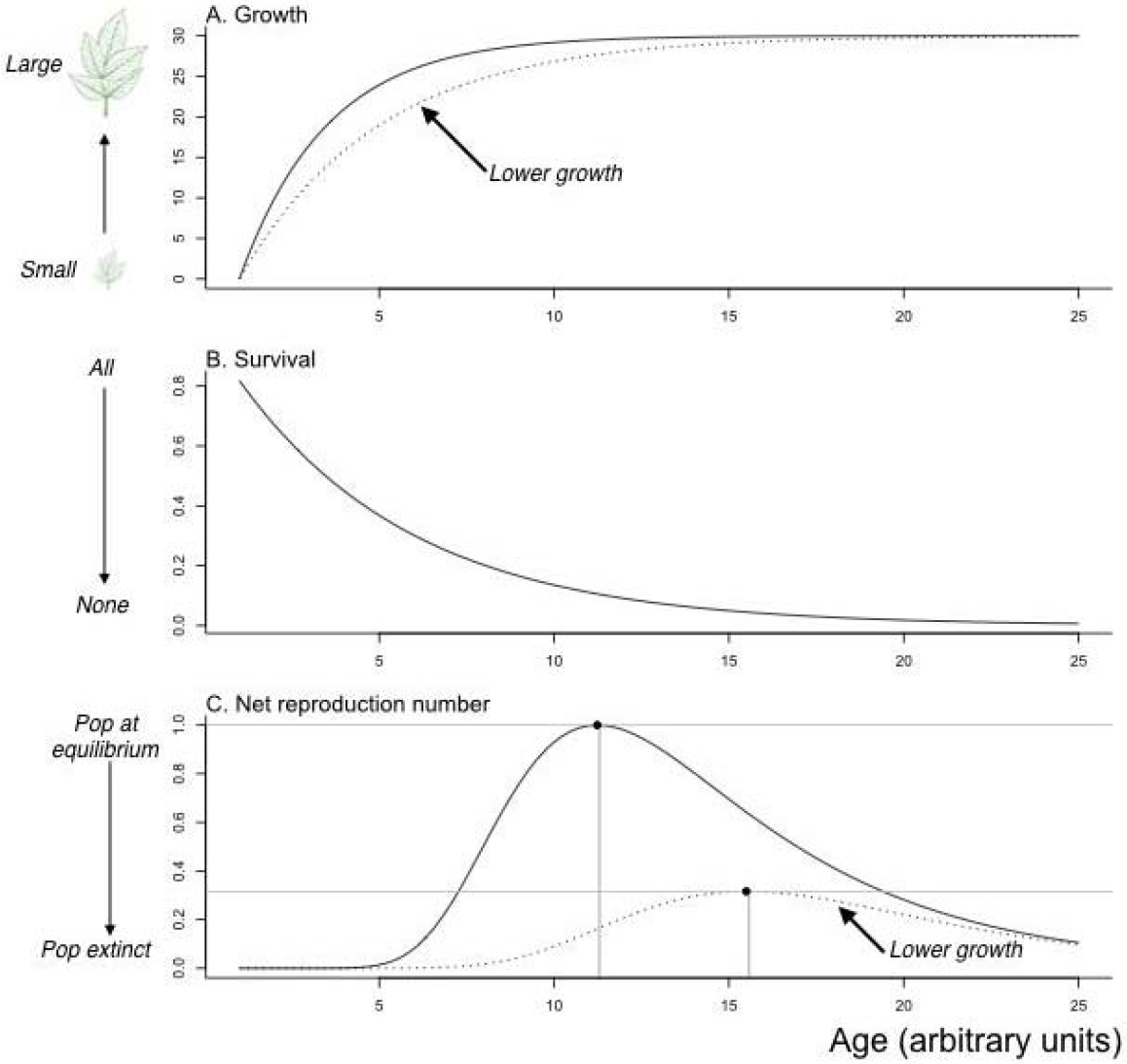
**Optimal timing of monocarpic reproduction** derived from a life history model where A) host individual growth in size (y axis) saturates with age (x axis) following *L*(*t*) = *L_T_*(1 − *exp*(−*k*(*t* − *t*_0_)))where *L_T_* is the maximum possible size, *k* defines the growth rate, and *t*_0_ is the hypothetical age at which size would be zero (here, *L_T_* = 30, *t*_0_ = land *k* = 0.4 (solid line), or *k* = 0.25 (dashed line)); B) Mortality occurs at rate *d*_0_, so that the probability of surviving until time *t* is *exp*(−*d*_0_*t*) (here *d*_0_ = 0.2). C) Combining these relationships with an expression for reproductive allometries (size is converted into offspring according to *S_i_* = *exp*(*A + BL*(*t*)); here, *A* = −5 and *B* = 1); and a probability of offspring establishment (here, *p_e_* = 2*e*^−10^ chosen to set one of the two populations at equilibrium), we can obtain an expression for the net reproduction number, *R*_0_: since reproduction is fatal, *R*_0_, is defined by the number of offspring produced by an individual at its age of reproduction, t, *R*_0_ = *p_e_ exp*(−*d*_0_*t*)*exp*[*A + BL_T_*(1 − *exp*(−*k*(*t* − *t*_0_))))]. To identify the age at reproduction that maximizes fitness as measured by *R*_0_, we solve for *dR*_0_/*dt* = 0, which yields *t_opt_* = *t*_0_ + *log*[*kBL_T_/d*_0_]/*k* (vertical lines, here 11.23 And 15.49); see supplementary code.

**Table 2:**
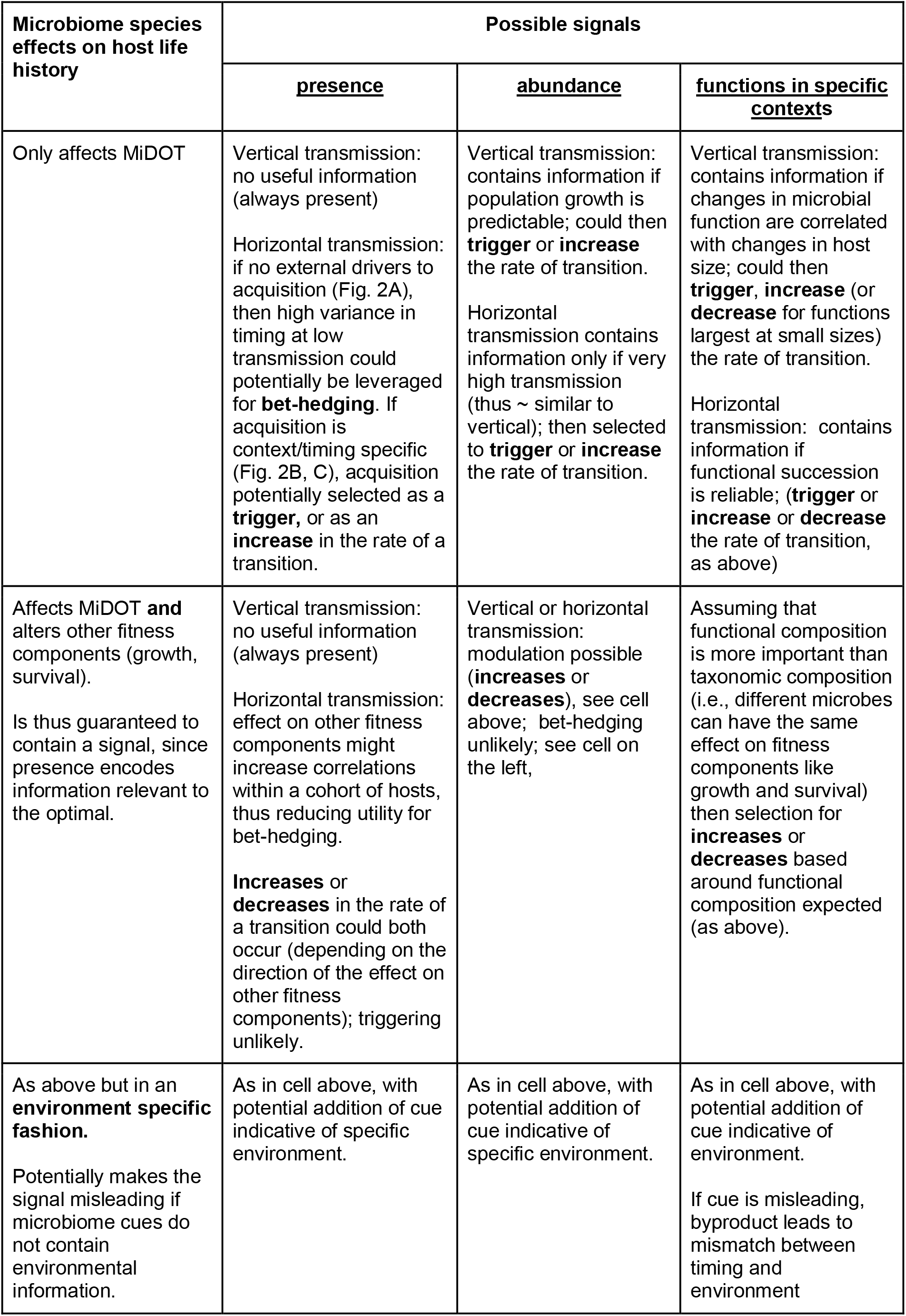
<u>M</u>icrobiome <u>D</u>ependent <u>O</u>ntogenetic <u>T</u>iming (MiDOT) in a life history context. Categorizing MiDOT via its effects across the life history (first column), and the information encoded by presence / abundance / functions and byproducts (columns 2-4), for vertical or horizontal transmission. We focus on the example of a monocarpic species, and evaluate potential contributions to **optimizing timing** (either as a **trigger**, or as **in/decrease** in the rate of a transition, Figure 1) or **bet-hedging** (see text).

While much of this research has focussed on optimal timing (i.e., at what age or size a transition should occur), there is also a rich set of results relating to selection for variance in timing. If individuals are selected to hedge their bets across time-varying environmental conditions [17,18]; or varying timing can reduce crowding and therefore competition, thus increasing fitness [19,20], the optimal strategy requires individuals with the same genotype in the same environment to produce different phenotypes (intra-genotypic variance). Seed dormancy is perhaps the best understood example of a bet-hedging trait [14]; the mechanism underlying intra-genotypic variation is thought to involve micro-gradients across seeds [21], but this does not exclude a potential role for MiDOT.

Overall, both theoretical [12,22] and empirical [18,23,24] research indicates that the timing of life history transitions have important fitness consequences. Therefore, factors which accurately reflect appropriate life-history timing have the potential to be harnessed as cues for ontogenetic transitions. The outcome of such selection could be MiDOT. The next questions arising to understand the selective context underlying MiDOT are therefore clear: what determines the timing of acquisition of species in the host-associated microbiome during ontogeny? Are there expectations for patterns of timing of abundance of functionality that could similarly be leveraged?

### Determinants of the timing of microbiome acquisition, abundance and functionality

Microbiota acquisition may be vertical (transferred from parent to offspring), horizontal (microbial colonization after germination/birth/hatching from the environment or conspecifics) or a combination of both [25]. Acquisition of vertically transmitted microbiome species will be highly predictable (all offspring acquire their microbiome early in life). The presence of vertically transmitted bacteria will thus manifest little variation that could be harnessed to structure subsequent host developmental transitions. However, changes in population growth or bacterial phenotype after initial colonization (in response to environmental cues or host size) could generate the varying time-signature (or context-signature) required for MiDOT to be adaptive. Thus, under vertical transmission, we would expect the cue from microbial species involved in MiDOT to reflect some feature of microbial ecology, rather than microbial species’ presence alone. For example, upregulation of a microbial signalling molecule might act as a predictable signal to trigger a host developmental transition.

Considerably more variability is expected in timing of colonization for horizontally transmitted microbiome species, echoing the well-understood process of transmission for pathogens [26]. In infectious disease biology, the ‘force of infection’, or rate at which uninfected individuals become infected [27], is defined by the product of the prevalence of infectious individuals in the population, and the transmissibility of the pathogen (constant in the simplest case, but could also be modulated by host age, or climatic conditions, etc).

Applying the logic of the ‘force of infection’ to thinking about horizontal acquisition of microbiota delineates clear expectations for patterns of age- (or time-) incidence associated with particular horizontally transmitted microbiome species. If the ‘force of infection’, or rate of acquisition of a particular microbial species is a constant, *λ*, the probability of being colonized by that microbial species is 1 − *exp*(−*λt*) and the average age of acquisition (or potentially timing in the year) is 1/*λ* (Figure 2A); this rate also encodes variance in microbiome species’ presence across individuals in the population, defined by 1/*λ*^2^.

**Figure 2:**
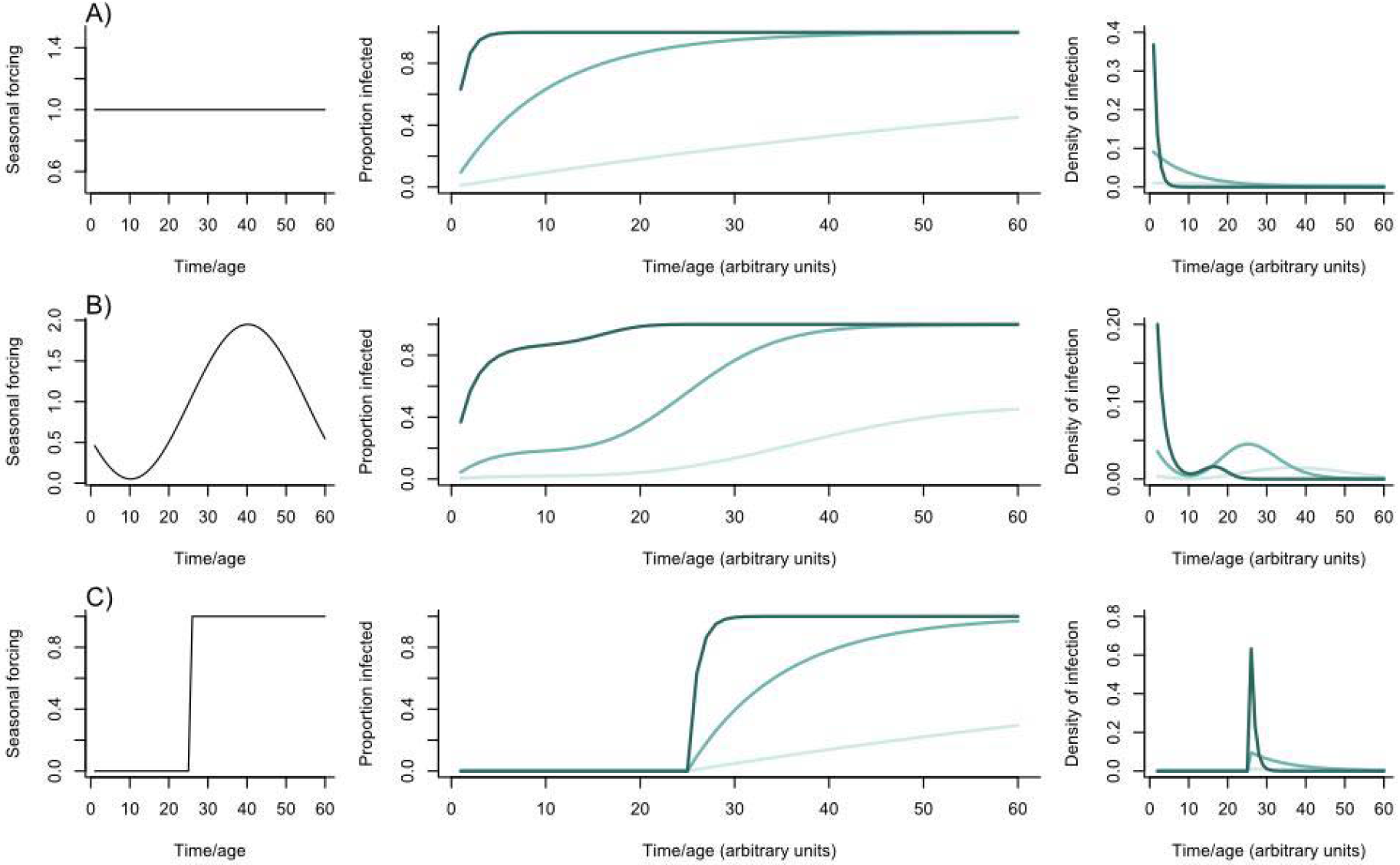
Timing information from the microbiome. For three magnitudes of the force of infection’, or rate at which susceptible individuals are colonized by species from the microbiome (*λ* = 0.01, *λ* = 0.1, and *λ* = 1, coloured from light to dark green, respectively), three different profiles of individuals being infected as a function of time (e.g., time during the year, or age) are obtained (second column), resulting in different patterns of age (or time) at infection (third column), with increased variance for lower forces of infection. The basic patterns shown in A) can be modulated by B) seasonal or C) abrupt changes in the force of infection, which can result in more or less narrowly defined age (/timings) of infection (last column).

Seasonal fluctuations in the force of infection represent another key way in which a microbiome species’ presence could be used as a host cue (Figure 2B,C). If aspects of the host affect microbiome acquisition (e.g., host behaviour, body size [28]), microbiome presence could act as a predictable indicator of host age or season/timing (noting that a high rate of transmission at the appropriate stage would be required to make MiDOT reliable). Beyond the temporal or context signatures indicated above, non-linear feedbacks inherent in transmission dynamics could add interesting scaling of information about population dynamics - more rapid acquisition might indicate higher density of conspecifics, and this potentially reflects information that is relatively inaccessible to the host otherwise (see below).

### When might microbes provide the best signal available to time transitions?

When might the microbiome provide the ‘best’ signal by which hosts can calibrate ontogeny? We first focus on the situation where there is a clear optimal timing, and evaluate three scenarios.

The first scenario is that presence (/abundance or function) of particular microbial species reflects the most accurate signal of either the hosts’ endogenous or exogenous state in terms relevant for optimally timing an ontogenetic transition. Many environmental conditions are likely to be closely mapped by microbial species’ presence or abundance [29], given their tight ecological dependence on environmental context (e.g., pH [30], temperature [31], drought [32]). For the example of drought, there also exists the intriguing possibility that microbial species can provide nuanced information as to past environmental conditions [33], which may also be pertinent information for timing of ontogeny - legacy effects within microbial community dynamics might mean that their abundance integrates over the longer term. For example, microbes might provide a better signal of drought than soil moisture if an overall dry season is interrupted by sporadic intense rain. Although microbial abundances vary unpredictably through time, microbial function might still be sufficiently linked to environmental conditions [34,35] to provide useful information. Finally, one feature of hosts’ ecology which may be key to the optimal timing of ontogenetic transitions is the presence and abundance of competitors.

Uniquely, species of the microbiome have the potential to provide information about this [36–38], because the ‘force of infection’ that defines horizontal transmission (and thus the rate of acquisition of species of the microbiome) is defined by the local abundance of hosts colonized by these microbiome species.

The second scenario is one where there are other, more accurate signals of the relevant endogenous or exogenous state; but these are not detectable by hosts. For example, it has been suggested that plants cannot readily identify their size, despite strong selection for size-based timing of flowering [39]. If microbe acquisition (/abundance or function) maps onto the relevant timing, hosts might be selected to respond to accessible microbial cues. In general, selection to respond to a cue will depend on the quality of the cue (how reliably it predicts changes in relevant endogenous or exogenous states), but also the difficulty or cost of acquiring the cue. In many cases, even though microbes might not be the best possible source of information, they might be the most detectable: animal and plant hosts have evolved multiple receptors and regulation pathways to respond to microbial activity [40] and these can be easily co-opted for ontogenetic regulation. Thus, microbes might provide reasonable information that is easy to detect.

The third scenario reflects situations where microbes modulate the conditions that shape optimal timing of flowering (e.g., by altering the growth rate, Figure 1). As a result, microbe species’ presence and/or abundance provides direct information as to how timing of an ontogenetic transition might be modulated to maximize fitness [3]. For example in a monocarpic plant, the presence of a species that accelerates growth indicates that early flowering will be associated with higher fitness (Figure 1C), all else being equal.

To illustrate how life history theory and microbiome ecology can be combined to develop a predictive framework, we lay out expectations for selection on MiDOT (Table 2), for the example of timing of reproduction in a monocarpic species (Figure 1). Here, optimal timing is driven by endogenous factors, i.e., plant size (rather than exogenous factors like seasonal context), and we assume for simplicity that acquisition and abundance of the microbiome tend to increase monotonically with host age or size (but see [41]). Organizing the ecological drivers in the context of known evolutionary selection pressures in this way (Table 2) yields clear predictions: i) unless the microbiome species affect a host trait other than MiDOT, only ‘triggering’ or ‘increases’ in the rate of the ontogenetic transition are expected; ii) MiDOT is most likely to be associated with bet-hedging in scenarios where microbiome species only affect MiDOT; iii) if microbiome species affect host growth, the direction of the effect will allow prediction of the expected effect of MiDOT. With more detail on particular microbiome species, or knowledge of key ontogenetic cues, such predictions could be made more specific, opening the way to formal tests of the selective value of MiDOT.

### Future directions

Characterizing if and when MiDOT is adaptive (Table 2), has potential applications from increasing agricultural yield to pathogen control. This line of research might also provide a broader physiological perspective on when presence of particular microbiome species is more tractable and accessible to the host than direct measurement of endogenous and exogenous determinants of optimal timing, such as host size [28], or environmental conditions [18].

When the benefits of using microbiome cues outweigh the costs of potential exploitation emerging from misaligned incentives is a question ripe for detailed investigation. Taking the example of monocarpic species, delaying reproduction is likely to be desirable for any species that live on or in a host (unless vertically transmitted); increased growth in size through delayed reproduction is also likely to benefit species for whom hosts are habitat. Host manipulation by microbes for delayed reproduction then seems likely, and even tractable (e.g., where microbiome species modulate insulin pathways [2]). Alternatively, for species of the microbiota which are directly consumed by the host, accelerating development to reduce host fertility and life expectancy (as observed in *C. elegans* [42]) might be optimal. What determines the trajectories of such co-evolutionary pressures? Are within-host within-microbiome interactions sufficiently uncoordinated and competitive that they are unlikely to drive host outcomes in directions that can optimize microbiome fitness [43]? Are there situations where this is not the case? Bounding the set of contexts where MiDOT occurs and could be adaptive will open the way to evaluating the relative importance of co-option of host ontogeny.

Identifying tractable systems for probing these questions is an important step. Known host microbiome alliances (Table 1) are likely to be key. Simpler systems, including synthetic microbiomes of model systems, may also provide important insight. In the closest living relatives of animals, both the transition to multicellularity, and the adoption of sexual reproduction, hinge on molecules produced by bacteria in the environment, highlighting how widespread such dependencies are, and providing an important potential test-bed for evolutionary predictions [29]. Conversely, our overview provides few examples of MiDOT in vertebrates species, potentially as a result of more complex buffering by the immune system (germ-free vertebrates often experience developmental issues associated with immunity [44]); but it might also simply be that more data is required. Overall, the increasingly detailed resolution available on microbiome species’ effects on the biology of a diversity of hosts provides the exciting opportunity of probing the broad phylogenetic context of MiDOT, and using theory and data to robustly evaluate the degree to which this surprising phenomenon is adaptive.

## Supporting information

R code for Figures

